# Measuring fingerpad deformation during active object manipulation

**DOI:** 10.1101/2021.07.20.453037

**Authors:** Benoit P. Delhaye, Félicien Schiltz, Allan Barrea, Jean-Louis Thonnard, Philippe Lefèvre

**Affiliations:** Institute of Neuroscience, Université catholique de Louvain, Brussels, Belgium; Institute of Information and Communication Technologies, Electronics and Applied Mathematics, Université catholique de Louvain, Louvain-la-Neuve, Belgium

## Abstract

During active object manipulation, the finger-object interactions give rise to complex fingertip skin deformations. These deformations are in turn encoded by the local tactile afferents and provide rich and behaviorally relevant information to the central nervous system. Most of the work studying the mechanical response of the finger to dynamic loading has been performed under a passive setup, thereby precisely controlling the kinematics or the dynamics of the loading. However, to identify aspects of the deformations that are relevant to online control during object manipulation, it is desirable to measure the skin response in an active setup. To that end, we developed a device that allows us to monitor finger forces, skin deformations, and kinematics during fine manipulation. We describe the device in detail and test it to precisely describe how the fingertip skin in contact with the object deforms during a simple vertical oscillation task. We show that the level of grip force directly influences the fingerpad skin strains and that the strain rates are substantial during active manipulation (norm up to 100%/s). The developed setup will enable us to causally relate sensory information, i.e. skin deformation, to online control, i.e. grip force adjustment, in future studies.

## Introduction

Humans are exquisitely skillful at dexterous manipulation of objects with their fingers. This dexterity partly relies on optimally adjusting the grip force (GF), the force exerted normally to the object surface, to the load force (LF) due to the object weight and inertia, and the fingertip-object contact friction (Johansson and Flanagan 2009a). Indeed, an excessive amount of force is undesirable as it leads to excessive expenditure of energy and could potentially crush the object. However, an insufficient grip force will let the object slip from the hands. Accordingly, we usually exert a grip force just above the minimum, thereby continuously varying the GF according to the LF and the level of friction (Cadoret and Smith 1996; Flanagan and Wing 1997; Johansson and Westling 1984; Westling and Johansson 1984). Deformations and vibrations produced in the skin during object manipulation are faithfully encoded by the numerous tactile afferents innervating the hand and fingers. These afferents in turn provide the central nervous system with rich and behaviorally relevant tactile information content (Delhaye et al. 2018, 2021; Goodwin and Wheat 2004; Jenmalm et al. 2003; Macefield et al. 1996). This, for instance, allows the central nervous system to adjust the grip force exerted on a manipulated object to the grip conditions (Johansson and Flanagan 2009b). Without tactile afferent feedback, we fail to react to unexpected perturbations in object load or surface friction and to maintain a stable grip force to load force ratio, hence highlighting the critical role played by tactile feedback during object manipulation (Augurelle et al. 2003; Nowak et al. 2001; Witney et al. 2004).

In vivo biomechanics of the fingerpad has been extensively studied in passive conditions, highlighting the systematic occurrence of partial slips at the periphery of the fingertip-object contact during the tangential loading of the fingerpad (André et al. 2011; Delhaye et al. 2014, 2016; Tada et al. 2006; Tada and Kanade 2004). Indeed, we have shown that slip is not instantaneous and that the transition from a stable to a slipping contact develops progressively with partial slips initiating at the periphery and progressing towards the center of contact until the point of a full slip. As a consequence of the relative movements between the non-slipping regions and the slipping regions, partial slip is associated with substantial (up to 25%) surface-tangential skin strains (Delhaye et al. 2016). Furthermore, we have shown that those skin strains caused by partial slips are readily encoded by human afferents (Delhaye et al. 2021). Besides, we have also demonstrated that human subjects can consciously perceive incipient slip, before full slip is reached, given that the surface-tangential skin strains are sufficient (Barrea et al. 2018). Taken together, those results have essential implications for object manipulation: indeed, the systematic occurrence of partial slips during tangential loading implies that partial slips will take place during active manipulation. Because partial slips provide a measure of how far from fully slipping the contact is, they might be used by the central nervous system to adjust the GF to the object properties during active manipulation.

It remains unclear, however, how much partial slips spread inside the contact area during active object manipulation and how they will be affected by the gripping conditions, including the biomechanical properties of the fingertips. Indeed, for a given object load, the amount of partial slips depends on how much grip force is exerted on the object and also on the frictional properties of the fingertip-object contact (André et al. 2011).

To investigate this, we developed a new instrumented device that enables synchronously recording the forces exerted by the fingers together with the fingertip skin deformation at the finger-object contact with a high spatial and temporal resolution during active manipulation. The device was tested in an experiment involving 18 subjects who were requested to perform vertical oscillatory movements. We describe the typical strain patterns taking place in the contact in parallel with the kinematic and dynamic parameters of the task.

## Methods

### Apparatus

To characterize the deformations taking place at the contact between the fingerpad and an actively manipulated object, we developed a manipulandum equipped with force sensors as well as an imaging system. This system can capture images of the skin in contact with the object through a transparent and optically flat plate of glass (Figure 1).

**Figure 1:**
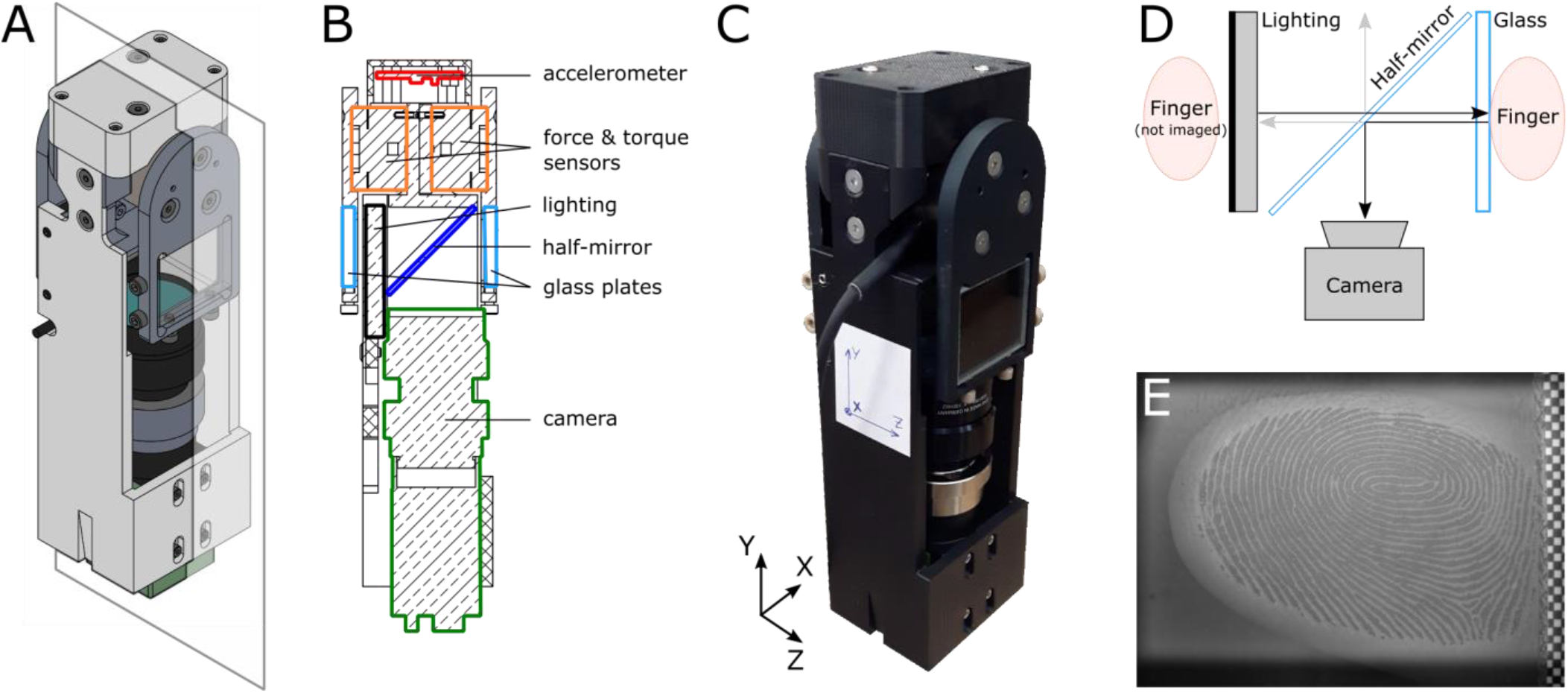
Detailed description of the device. **A|** Assembly sketch of the device. **B|** Cut of the device along the plane in panel A with its key elements and sensors highlighted. **C|** Photograph of the manipulandum along with the definition of the reference frame used in the context of this study. **D|** Custom optical system to image fingerprints. Arrows show the trajectory of the light emitted by the “Lighting” and reflected at optical interfaces. In this study, the index finger was imaged, and the thumb was not. **E|** Typical image obtained with the system. The checkerboard pattern on the right of the image is used to measure very small movements of the glass relative to the camera.

The device is designed to be handled using a precision grip (Westling and Johansson 1984), i.e. pinched between thumb and index. Its aluminum core supports two force sensors, one on each side of the manipulandum, to measure the forces exerted by each finger (Fig 1B, orange, ATI Mini27 Ti, ATI-IA). On the outer side of each force sensor, an aluminum piece supports a transparent plate of glass, where the object is contacted to be manipulated. The rest of the device, including its top hat and diverse small parts, is 3D printed using PLA to be lightweight and customized at will (Fig 1A-C). Only one finger can be imaged, but the outer plate design is symmetric so that the contacting surfaces are the same for both fingers. Both surfaces can be removed and swapped quickly, allowing the experimenter to test materials of different properties (e.g. friction or texture) during one experiment. An accelerometer (Fig 1B, red, LIS344ALH, STMicroelectronics) is included in the top hat of the device. All signals are acquired using an ADC acquisition system (NI6225, National Instruments).

Fingerprints in contact with the glass plate are imaged using a custom optical system based on the principle of frustrated total internal reflection, with a coaxial light source and camera (Fig 1D). The light source (Fig 1B, black, LFL-1012-SW2, CCS Inc.) is placed on the opposite side of the imaged finger. The light passes through a half-mirror to illuminate the finger. Part of the light is reflected by the contacting glass and part of it is transmitted. The light transmission index is increased where the fingerprints contact the glass, causing less reflection and therefore dark fingerprints in the captured images (Fig 1E). The light is then collected by a small and lightweight monochrome camera equipped with a macro lens (Fig 1B, green, camera: GO-5000M-PMCL, JAI, monochrome, 2560 × 2048 full pixel resolution; macro lens: Cinegon 1.8/16, Schneider Optics). The setup enables us to image the entire fingerprint at a high spatial (1696 × 1248 pixels) and temporal (100 Hz) resolution. Note that only a subset of the camera field of view was recorded because of constraints related to the architecture of the device, in particular the size of the area illuminated by the light source. Optical tracking allows us to measure partial slips from fingerprint images and to derive the skin strains (Delhaye et al. 2014, 2016). A checkerboard pattern is glued on the glass on the side of the field of view to track very small glass movements relative to the camera caused by the elastic deformation of the device. The squares of the checkerboard, whose dimensions are 0.5 × 0.5 mm, enable us to measure the image resolution (64 pixels/mm) and to verify that the effects of optical distortion and elastic deformation of the manipulandum are minimal. The width of the device, i.e. the distance between both fingers gripping the object, is 50 mm. The total mass of the device is 540 g.

### Participants

Eighteen healthy subjects (9 women, ages 20-34 years) participated in the experiment. Each subject provided written informed consent to the procedures and the study was approved by the local ethics committee at the host institution (Institute of Neuroscience, Université catholique de Louvain, Brussels, Belgium).

### Experimental procedures

For this experiment, the device was held by a system of pulleys and a counter-weight with the same mass as the device, such that the net weight of the whole system was close to zero, but its inertia was twice that of the device alone. The goal was to isolate the forces and deformations related to dynamic interactions (inertial forces) as opposed to static weight. Moreover, an optical distance sensor (DT20-P224B, SICK Sensor Intelligence) was placed above the device and measured its vertical position continuously.

Subjects were asked to grab the device with their index and thumb at the center of the glass plates such that the fingerprints were centered in the field of view of the camera (Fig 1C-E). Then, they were instructed to move the device up and down producing an oscillatory movement between two targets spaced by 20 cm for 30 seconds (Figure 2). The period was set to 1.5 seconds, therefore generating 20 oscillations for each block (Fig 2A). A short tone was played every 750 ms to indicate the instant when they were supposed to reach one of the targets. A higher-pitched tone indicated the end of the block when the subjects were asked to release the object and prepare for the next trial. The same procedure was repeated for 15 blocks. Because we observed that subjects tended to apply an excessive level of grip force, a subset of them (6) were explicitly asked to try to minimize the grip force.

**Figure 2:**
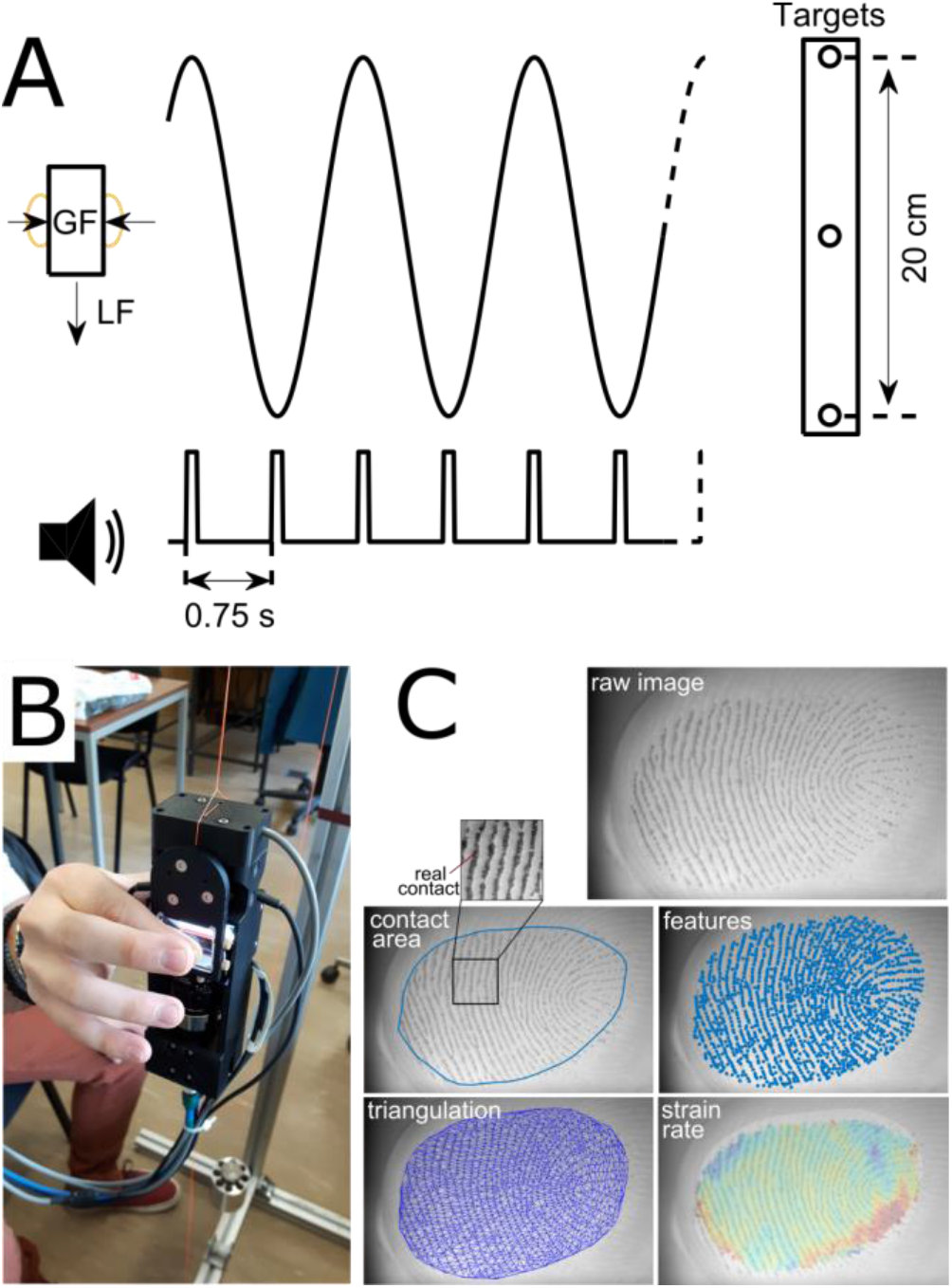
Experimental protocol and image processing. **A|** Subjects performed vertical oscillations with a peak-to-peak amplitude of 20 cm, delimited by visual targets and rhythmed by auditory cues every 0.75 s (at each peak of the movement). **B|** The device was held in a precision grip. A system of pulleys and a counter-weight completely compensated the weight of the device. **C|** Image processing pipeline. From the raw images, the contact area is extracted. Feature points are then detected inside this contact area and tracked from frame to frame. A Delaunay triangulation is computed from the first frame and the strain rate tensor of each triangle can be computed for each pair of images.

Moreover, at the end of the experiment, the subjects were asked to rub their index finger against the glass surface repeatedly using a published procedure (Barrea et al. 2016) to evaluate the coefficient of friction as a function of the normal force. A power function was fit to the data (*μ* = *k*(*NF*)^n-1^, where *μ* is the coefficient of friction). It was used to predict the slip force.

The forces, torques, position, and acceleration were acquired at 200 Hz. The images were captured at 100 frames per second at a resolution of 1696 by 1248 pixels.

### Data analysis

Force, torque, and position signals were low-pass filtered (4^th^ order Butterworth, zero phase-lag, cutoff 40Hz). The grip force (GF) was obtained by computing the average of the force normal to the surface from the two force sensors (along the z-axis, see Fig 1C). The load force (LF) is the sum of the vertical component of the tangential force from the two force sensors (along the y-axis, see Fig 1C). The object velocity was obtained by numerical differentiation, and the positive velocity peaks were used to split the data into individual oscillations during which different parameters were computed (mean GF, max LF,…). The vertical position of the center of pressure (COP) was obtained using *COP_y_* = (*T_x_* — *F_y_*. *z*_0_)/*F_z_*, where T is torque along a given axis, F is force and *z*_0_ is the normal distance between the contact surface and the sensor’s surface. It measures the vertical position of the resultant force exerted by the index finger normally to the glass. The slip force (SF) is the minimum normal (grip) force needed to avoid slip. It was obtained from the power function fit for each subject as a function of the tangential force (*TF* = *μ*.*SF* = *k*(*SF*)^*n*^). The GF and LF modulations were obtained for each oscillation by subtracting the minimum value from the maximum value of the specific oscillation.

### Image processing

We used an image processing pipeline already described in previous work to detect the slipping regions and evaluate the surface skin strains inside the contact area (Delhaye et al. 2014, 2016). It is summarized here (see Fig 2C). First, the gross contact area was extracted from the background using a two-stage procedure. The first stage consists in manually depicting the contact area contour for a small subset of frames (typically 10 to 20 frames per subject). In the second stage, features relevant for the segmentation (Sankaran et al. 2017) are computed and fed to a machine learning algorithm (classification tree, *fitctree* function in Matlab) that is trained on the manually segmented data to extract the contact area and then used to extract the contact area on the whole sequence of frames.

Second, for each oscillation, three sets of features having good gradient properties for tracking (Shi and Tomasi 1994) were sampled at the first, the middle, and the last frame of each oscillation respectively. Indeed, the contact area varies during the movement due to rolling of the fingertip, i.e. some skin parts are coming into contact and other parts are leaving contact during the active movement. It is therefore essential to re-sample features at different times to cover the contact area with features during the whole oscillation. The minimum spacing between features for feature detection was set to 17 pixels. Then, those features were tracked from frame to frame using a classical algorithm implemented in OpenCV (Bradski 2008; Lucas and Kanade 1981). Depending on the initial frame (first, middle or last), the features were either tracked forward or backward in time (or both). Finally, the 3 sets of features (first, middle and last) were merged, and those overlapping were suppressed. The very small glass movements that occurred due to the compliance of the force sensors and the plates (of the order of a couple of pixels) were monitored by tracking features located on a checkerboard attached to the glass. The velocity of the glass movement was subtracted from the velocity of the fingerprint movement before further analyses.

Third, a Delaunay triangulation was performed on the features from their location in the first frame. And the triangle strain rate from frame to frame was computed as described earlier (Delhaye et al. 2016), to yield a 2-by-2 strain rate tensor for each triangle and each pair of consecutive frames (Fig 2C). The strain rate norm was obtained by computing the norm of the strain rate tensor for each triangle and each pair of consecutive frames. Moreover, if the triangle’s center moved by more than 1/2 of a pixel between 2 consecutive frames, it was considered as slipping whereas if the movement was smaller than 1/2 pixel, it was considered non-slipping or stuck to the glass. The *stick ratio*, the ratio of non-slipping area to the total contact area, was obtained for each pair of consecutive frames.

### Statistical analyses

All statistical analyses were performed in MATLAB, using the functions *corr* (for Pearson correlation), *ttest* (for paired t-tests) and *regress* (for linear regression). The power-law fits were obtained by computing the coefficients of a linear regression on the logarithmic transformation of the data.

## Results

We asked subjects to perform paced vertical oscillations holding the manipulandum in precision grip, i.e. with the thumb and index finger (Fig 2B). Figure 3 shows the evolution of the recorded variables during a typical experimental block. A video of the fingerpad during a typical trial is also provided in the supplementary materials. The manipulandum weight was fully compensated by a counterweight, and therefore LF was only due to the object acceleration and was zero when the object was still. Thus, as expected, the manipulandum movement (Fig 3A) generated cyclic fluctuations in LF due to the inertial forces, and we observed two peaks of load force that were very similar in amplitude, one in the upper and one in the lower part of the trajectory. The load force variations were accompanied by variations of the grip force applied by the subjects (median correlation across oscillations, r=0.61±0.17, mean±std across subjects, n=18, see also herebelow and Figure 6), and LF peaks gave rise to synchronized grip force peaks (Fig 3B). These GF peaks were such that the GF/LF ratio was kept above the friction limit in most cases (in red, Fig 3C), and therefore the object did not slip. However, even though full slip was avoided during most oscillations, the zone of stable contact also fluctuated, as partial slip was observed and was quantified by the stick ratio (SR, Fig 3D), which typically reached a minimum when the constraints were maximal, that is at the maximum of LF or the minimum GF to LF ratio. As described earlier in a passive context (Delhaye et al. 2016), partial slip is accompanied by significant strain patterns in the slipping regions. The heatmaps provided in the bottom part of Figure 3 show the evolution of the three independent components of the strain rate tensor at each point of the contact area during one oscillation (Fig 3E). The last line shows the norm of the strain rate tensor (Fig 3F, see *Methods*). While it does not bear any physical meaning, the strain rate norm provides a clear picture of the distribution of the strains and their intensities inside the contact area, irrespective of the type of deformation and invariant to rotation. As expected from previous reports, the strains take place at the periphery of the contact and propagate further toward the center as the SR decreases, i.e. when the GF to LF ratio comes closer to the friction limit. As shown in a typical trace in Figure 3, each oscillation was isolated based on its trajectory (Fig 3A, see *Methods*) and analyzed separately.

**Figure 3:**
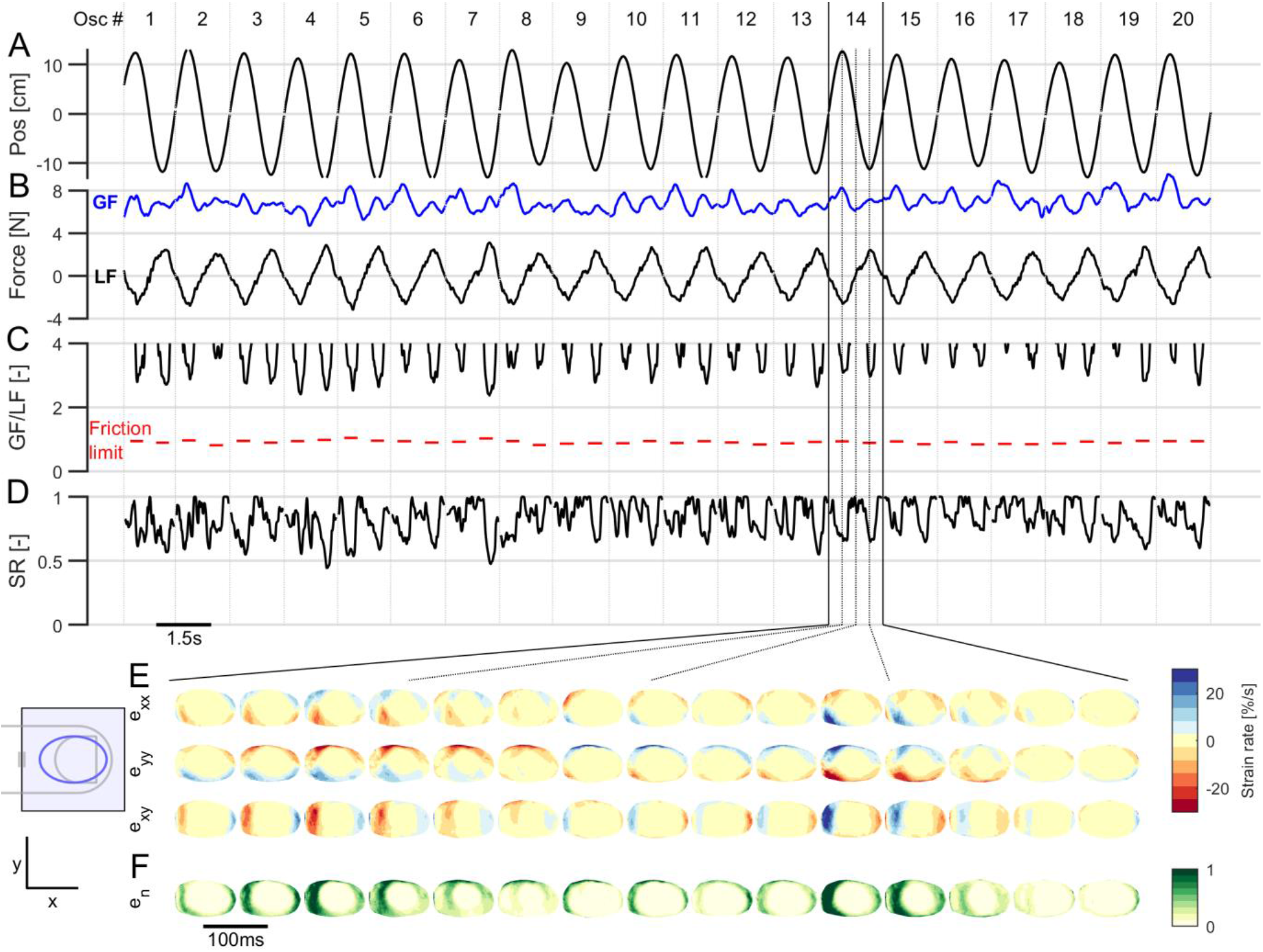
Typical trace. **A,B,C,D|** Evolution of the manipulandum vertical position (**A|**), fingertip forces (grip force, in blue, and load force, in black, **B|**), slip margins (grip force to load force ratio, in black, and friction limit at the load force peak, in red, **C|**) and stick ratio (**D|**) as a function of time during a typical experimental block. The oscillations are delimited with vertical lines at the times of positive velocity peaks. **E,F|** Evolution of the strain rate tensors, shown as three heatmaps for e_xx, e_yy, and e_xy (**E|**), and the strain rate norm (normalized by the 90^th^ percentile, **F|**) as a function of time during a typical oscillation. The sketch of the finger on the left shows the orientation of the finger with respect to the heatmaps (strains are shown as if viewed through the finger). The red parts are negative and show compression, and the blue parts are positive and show stretch. Consecutive heat maps are separated by 100 ms.

### Forces and strain patterns during oscillations

Since all signals followed a stereotyped pattern for each oscillation, we looked at the averaged evolution of the traces during all oscillations (Figure 4). The GF, LF, and SR patterns follow the behavior described earlier (Fig 4A-B, see also Figure 3). That is, the maximum GF and the minimum SR were observed on average at the time of the maximum LF. We also looked at the vertical displacement of the center of pressure (COP, Fig 4C), which is mainly caused by a redistribution of the pressure inside the contact area and the rolling of the finger. We observed that the COP moved significantly downward during the first half of the oscillation and upward during the second half of the oscillation. Even though each oscillation showed a slightly different pattern of deformation and widely different levels of minimum SR across subjects (see below), the general shape follows a trend that is expected from previous work (Delhaye et al 2016). The strain rate amplitude, as measured by the 90^th^ percentile of the strain rate norm across the entire contact, typically followed a trajectory similar to the variation of LF, or LF rate. Fig 4E shows the averaged pattern of strain rate during one oscillation for a typical subject. Since the central stuck zone is pulled up by the glass during the upper part of the trajectory (and pulled down during the lower part of the trajectory), the vertical strains (e_yy) were tensile in the lower regions of the contact area and compressive in the upper region of the contact area (Fig 4E). As a consequence of the elastic properties of the skin, horizontal strains (e_xx) appeared following the same pattern as the vertical ones, except that they were tensile where the vertical strains were compressive and vice versa. They were also smaller in amplitude. Finally, shear strains were substantial on the proximal (left) and distal (right) parts of the contact area. The norm of the strain rate (e_n) shows the areas of the skin where the strains are the largest, irrespective of the direction of the strains. It can be observed from those that a wave of strains progresses from the exterior of the contact area towards its center as the stick ratio decreases. In Figure 3, the wave of strains doesn’t reach the center of the fingertip as the stick ratio doesn’t reach zero.

**Figure 4:**
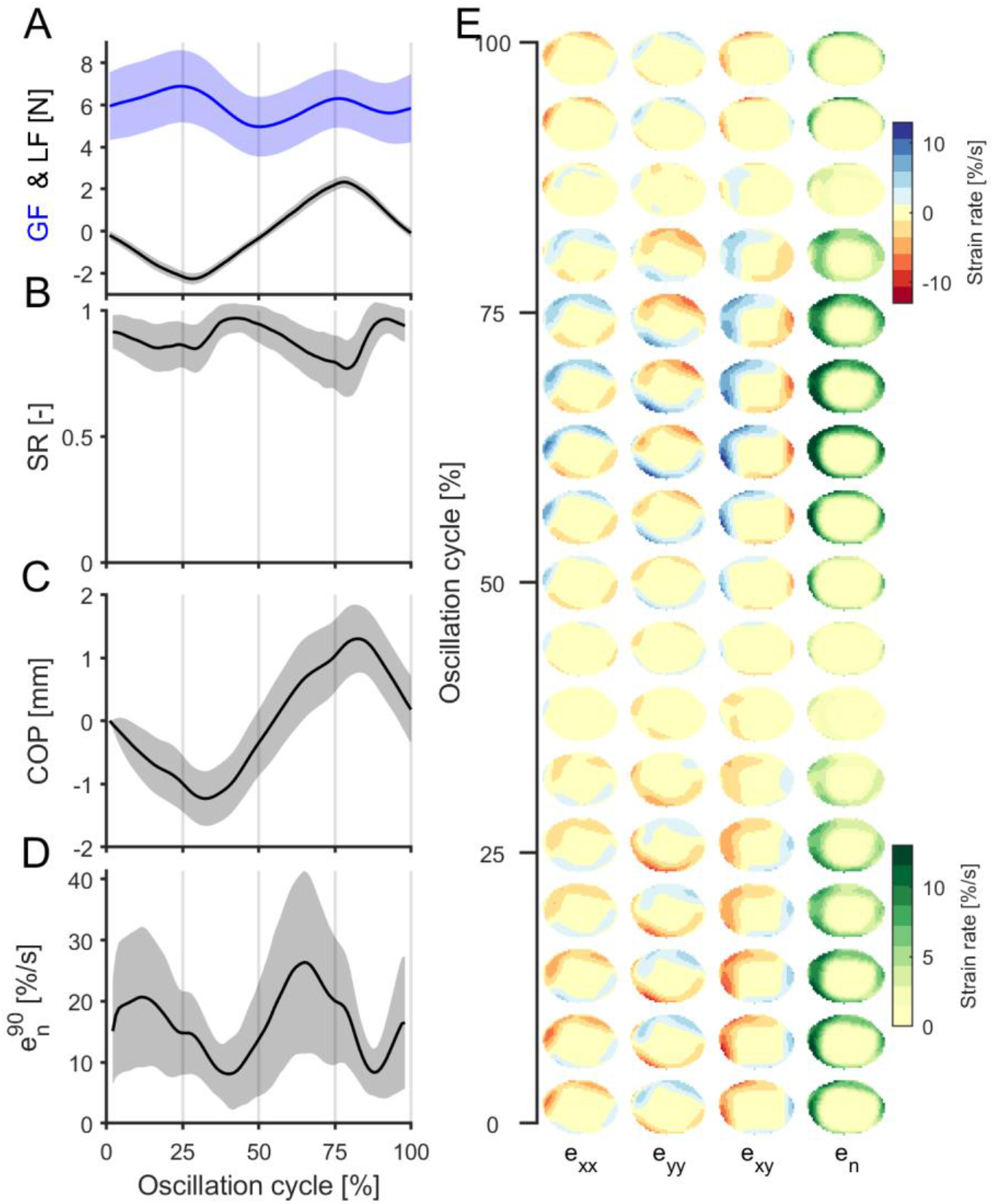
Oscillation averaged traces and strain rate patterns for a typical subject. **A,B,C,D|** Average traces across all oscillations expressed as a function of oscillation cycle progress (in percent) for a typical subject. The forces (grip force, in blue, and load force, in black, **A|**), stick ratio (SR, **B|**), the center of pressure (COP, **C|**), and 90^th^ percentile of the strain rate norm (**D|**) are shown. The shaded areas represent the standard deviations. The oscillations are cut using the positive velocity peaks and the signals are synchronized by expressing their evolutions in terms of percentage of time inside the oscillation cycle. **E|** Average strain rate patterns for a typical subject. Consecutive heat maps go from the bottom up and each column shows strain rates along a dimension. Strains were synchronized in the same way as the traces in A,B,C,D. Each strain heatmap is presented the same way as in Fig 3E-F.

### Different levels of grip force during manipulation

Looking at subjects’ gripping behavior (Figure 5), as described by the average GF value during one oscillation, we found that it was consistent across the entire experiment (Fig 5A). Indeed, even though the average grip force was on average slightly higher in the first block, this difference was not significant when compared to the last block (Fig 5C, paired t-test, t(17) = −1.14, p = 0.272). Within a block, there was a clear tendency of subjects to progressively decrease their grip force (Fig 5A). This was not related to a change in the movement kinematics, which should affect the load force (Fig 5B). Indeed, even though the first two oscillations of each block seemed to sometimes generate a higher load force (i.e. faster movement), the average load force of the third movement was already at the same level as the last movement (Fig 5D, right, paired t-test, t(17) = −1.59, p = 0.130). Interrestingly, GF progressively decreased during the experimental block and the difference between the grip force of the third and the last movement was highly significant (Fig 5D, left, paired t-test, t(17) = −3.67, p = 0.002), as previously observed during similar experiments (Augurelle et al. 2003).

**Figure 5:**
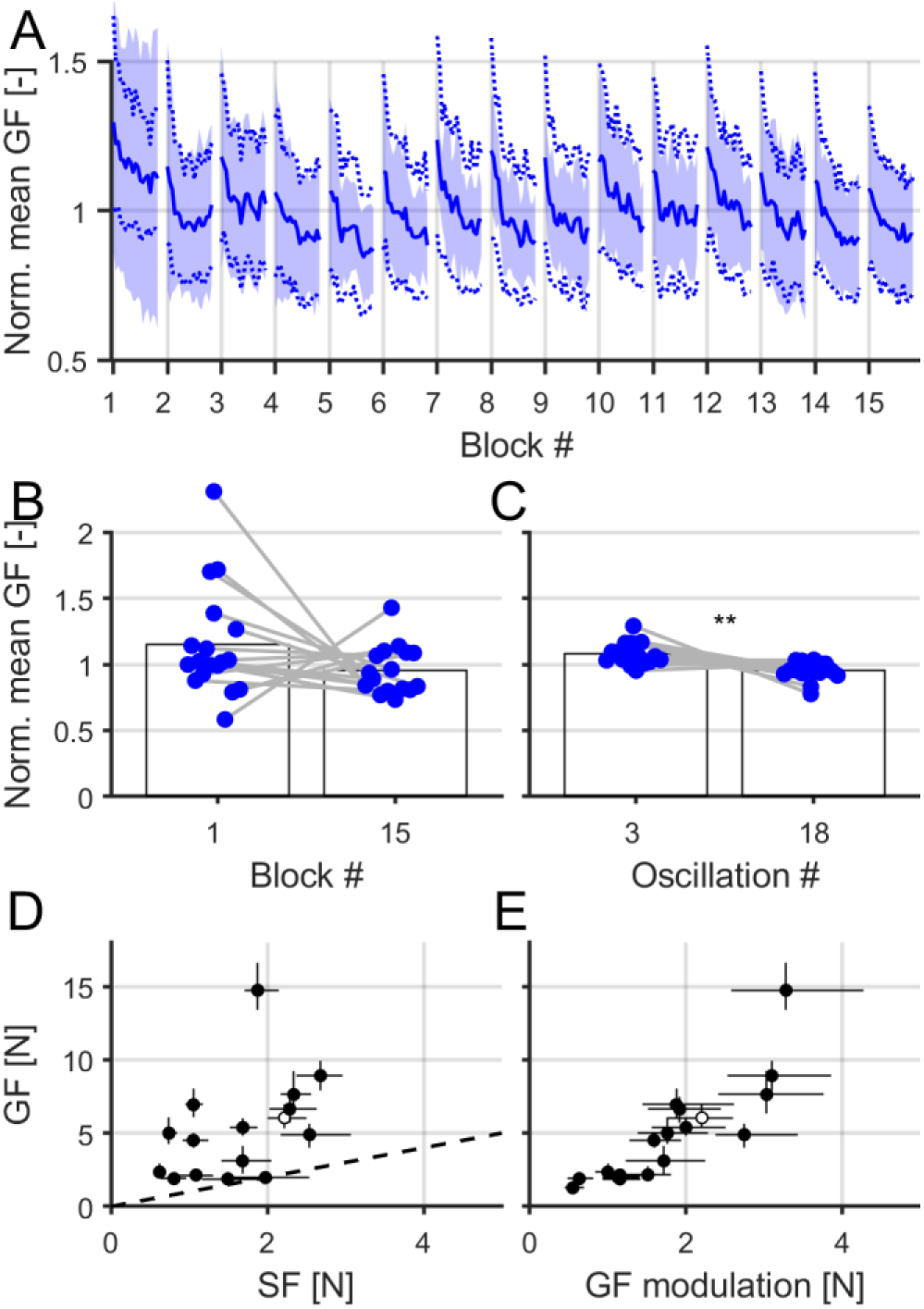
Gripping behavior. **A|** Evolution of the mean grip force (GF) across oscillations, blocks, and subjects (n=18). For each subject, the forces were normalized by the mean across all blocks and oscillations. Lines are mean across subjects and shaded areas show standard deviation. Dashed lines show means of minimum and maximum grip forces. **B|** Comparison of grip force at the first and last block (averaged across oscillations from those blocks). **C|** Comparison of grip force at the third and 18^th^ oscillation (averaged across blocks). Pair in B and C were t-tested, significant results are marked with stars (**: p<0.01). **D|** Grip force (GF) as a function of slip force (SF) as the instant of minimum GF to LF ratio. The slip force is the minimum grip force required to avoid slipping. The dashed line is the identity line. **E|** Grip force (GF, same as in E) as a function of the grip force modulation. In **D** and **E,** one point corresponds to the median value for one subject, the vertical and horizontal whiskers show the interquartile, the empty dot corresponds to the subject shown in Figure 4.

While the skin deformation patterns were similar across subjects and the grip force levels were constant across the experiment, we observed that different subjects used very different levels of grip force leading to very different amplitudes of deformations. Indeed, we found that the subjects that were instructed to use a minimal GF tended to apply an amount of GF just above the minimum required by friction, as shown by the peak in GF being just above the slip force estimated thanks to the friction measurements (Fig 5E). However, other subjects applied an excessive amount of GF. This is not resulting from the subject not coordinating their GF with the LF, as the subjects applying excessive GF also showed a high level of GF modulation, thereby taking into account the LF modulation (Fig 5F). It was rather explained by a high level of mean GF (Fig 5F).

### Mechanical parameters related to different grip force levels

The different levels of GF that individual subjects used to perform the oscillations led to different observations about the parameters of the manipulation (Figure 6). First, the vertical displacement of the COP (see *Methods*) tended to strongly decrease with the grip force level (Fig 6A). This decrease was observed across subjects, with a relationship that followed a negative power law (Fig 6A left, R^2^=0.91, F(1,16)=165.65, p<0.001, *f*(*x*) = *a*. *x^b^*, with a=10.93 and b=-0.84). It was also observed within-subjects, with a negative correlation that was significant at the group level (Fig 6A, right, paired t-test, t(17) = −4.45, p<0.001). This trend was not followed for the two subjects with the lower GF values, probably because of the very small range of GF (as shown by the very small horizontal wiskers for the two blue dots in Fig 6A right). The skin displacement range was also quantified. It measures the maximal vertical range of motion of a fingerprint feature inside the contact area and therefore quantifies how much the skin moves within the slipping regions (Fig 6B, left). This variable is strongly correlated with the maximum level of skin strain rate (shown in Fig 6C, correlation r=0.88). Both variables decreased with the grip force, according to a power-law (Fig 6B, left, displacement range: R^2^=0.63, F(1,16)=26.99, p<0.001, *f*(*x*) = *a*. *x^b^*, with a=1.79 and b=−1.27; Fig 6C, left, strain rate norm: R^2^=0.61, F(1,16)=25.50, p<0.001, *f*(*x*) = *a*. *x^b^*, with a=84.37 and b=−1.04), and the negative correlation within subjects was also observed (Fig 6B, right, displacement range: paired t-test, t(17) = −3.55, p = 0.002; Fig 6C, right, strain rate norm: paired t-test, t(17) = −3.32, p = 0.004). Again, the within-subjects correlation was lower for some subjects having small variations of GF across trials.

**Figure 6:**
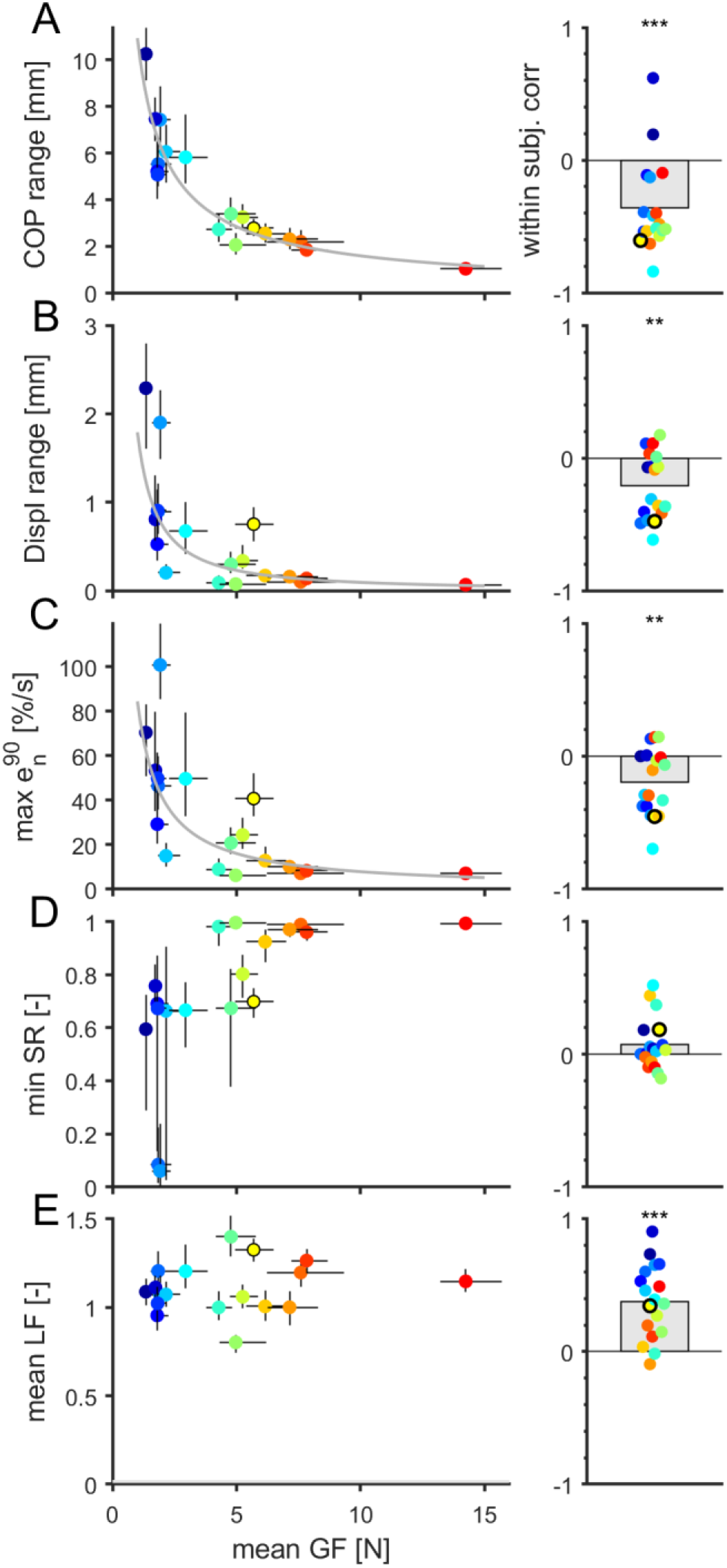
Mechanical parameters related to behavior. **A, B, C, D, E|** Relationship between different mechanical parameters and the grip force (GF), namely center of pressure range of motion (**A|**), fingerprint maximal range of motion (**B|**), maximal value of the 90^th^ percentile of the strain rate norm (**C|**), minimum of stick ratio (SR, **D|**), and mean load force (LF, **E|**). <u>Left panels</u> show the median across all oscillations of the different parameters as a function of mean grip force (GF) across oscillations for each subject. Dots are medians, vertical and horizontal whiskers show interquartile. <u>Right panels</u> show the intra-subject correlation of the parameters with the grip force, with the grey box indicating the average correlation across subjects. A t-test was performed for each variable and the stars depict the result of the test (nothing, p>0.05; *, p<0.05; **, p<0.01;***, p<0.001; n=18). Colors were attributed to each subject according to the mean GF level and correspond across panels. The dot with a black contour corresponds to the subject shown in Figure 4.

The maximal strain rates were substantial, and ranged from 6%/s to 100%/s (Fig 6D, left). Obviously, the maximal strain rates strongly correlated with the maximal strain values observed during an oscillation (r=0.99), which ranged from 1% to 17% depending on the subject’s GF level. Finally, as could be expected, the minimum value of the stick ratio within an oscillation was low for low levels of grip force and tended to 1 for very high levels of grip force (Fig 6D, left). We did not observe any significant tendency at the group level in the within-subjects correlation (Fig 6D, right, paired t-test, t(17) = 1.60, p = 0.128). As a control, we verified that the use of a high level of grip force was not related to the level of load force, but rather to an inappropriate adjustment to friction. We found that indeed, while there were some variations in the level of load force across subjects, related to variations in the movement acceleration from trial to trial, the correlation with the grip force was low (r=0.17, Fig 6E, left). However, within-subjects, we observed a clear positive correlation at the group level, meaning that the subjects adapted their level of grip force to the movement kinematics (Fig 6E, right, paired t-test, t(17) = 5.74, p<0.001). As could be expected, this tendency was strongest for the low GF values, for which the risk of slip is higher, and therefore a tight coupling between the LF and GF is required.

In summary, we showed that higher levels of GF yielded less fingertip rolling (as measured with the vertical displacement of the center of pressure), fewer skin strains (as measured with the strain rate norm) and a higher proportion of the skin remaining stuck on the glass (as measured with the stick ratio).

## Discussion

In this paper, we described a novel device that enables synchronous monitoring of strains in the fingertip skin in contact with a manipulated object and the forces exerted by the fingers on the object. The device was tested in an experiment involving 18 subjects and permitted to describe and quantify the strain patterns emerging from active object manipulation. This is the first study showing that substantial skin strain rates (>50%/s) take place inside the contact area during active manipulation, even though the object is in a stable, non-fully slipping contact and the object never dropped from the hand of the subjects.

The development of this device was inspired by a previous publication presenting a much simpler apparatus (Tada et al. 2002). This equipment was also manipulated in a precision grip and was also comprised of force sensors under each finger and of a camera to monitor the stick ratio, which relied on 167 manually drawn points on the fingertip to monitor skin displacement from frame to frame. Importantly, our setup has a much higher resolution (64 pixels per mm) and therefore enables an accurate measurement of local skin strains, which was not possible earlier.

Given the substantial amplitude of the strain rates measured in this study, it is reasonable to assume that those strains can be faithfully captured by the tactile afferents, and in particular the FA-I afferents (fast adapting type I afferents). Indeed, we recently demonstrated, in a passive setup, that FA-I afferents are sensitive to the compressive strain rates related to partial slip, comparable in scale to those observed here (Delhaye et al. 2021). Those strains therefore likely provide tactile feedback about grip safety.

There was a wide range of mean grip force levels used by different subjects observed in this study. Some subjects tended to apply a grip force level just above the friction limit while some others exerted a much higher grip force. Those behaviors lead to very different feedback. The “safe” strategy – exerting a high level of grip force – leads to a very limited amount of partial slip and a low level of deformation. Therefore, it also provides a limited amount of information related to contact stability. The “risky” strategy – using a grip force level just above the friction limit – leads to a large fraction of the contact area in partial slip and substantial strain rates inside the contact. Therefore, this strategy enables rich tactile feedback about the contact state at each oscillation.

While there was a clear relationship between the level of grip force and the amount of skin strains (Fig 6C), there was also a large variability. This variability can be explained by individual differences in skin properties. Indeed, the fingerpad can have very different mechanical and geometrical properties: stiffness (Wang and Hayward 2007), humidity (André et al. 2010), size (Peters et al. 2009), and all of those likely influence deformation and perception (Gueorguiev et al. 2016; Peters et al. 2009).

### Limitations

The developed device has some limitations. First, for a device that is intended to be used in precision grip, that is pinched between the thumb and index finger, its mass is rather large (around 500 grams). This limitation can be compensated by a counterweight, as done in this work, but it modified the weight/inertia relationship of the object, which makes it more remote from natural object manipulation. The manipulation of an object with no weight but high inertia, as done in the present study, is comparable to object manipulation in weightlessness in terms of contact forces with the object. Furthermore, due to the counterweight setup, we are limited to vertical movements. In addition, we used smooth transparent glass. While this is a very convenient material for imaging the skin-object contact, the flat contact with a glass surface is very different from the rough contact experienced with most natural objects.

As summarized above, some subjects tended to use a high level of grip force, much higher than the friction limit. This constrats with previous work showing that, under normal sensory feedback, people tend to manipulate objects with a grip force close to the slip limit (Augurelle et al. 2003). This might be explained by three factors. First, the instrumented object might look fragile and subjects tended to apply excessive GF levels to make sure not to drop it. Second, by compensating the weight of the object with a counterweight (see *Methods*), the experienced weight is much lower than the one expected from the object’s appearance and size, which may surprise the subjects. Moreover, there is a discrepancy between the perceived weight of the object, which is zero, and the interaction force resulting from the doubled inertia. As a result, it is likely that an excessive GF is exerted by caution. Finally, glass remains an material having particular frictional properties, with a coefficient of friction that can vary by up to one order of magnitude depending on several factors (Adams et al. 2013; Delhaye et al. 2014; Pasumarty et al. 2011), and might therefore encourage higher safety margin even if not needed.

### Conclusion

The device developed in this study will enable the monitoring of fingertip skin strains at the finger-object contact and provide a window into the feedback from tactile afferents inside the contact area during manipulation. Future work will test if unexpected changes in the object parameters, such as a change in friction, can be quickly detected and accounted for thanks to the feedback provided by partial slip.

## Supporting information

Images of a typical trial

## Acknowledgments

The authors thank Hélène Nokerman for collecting part of the data,François Wielant for his help throughout the development of the device and David Cordova Bulens for useful comments on a previous version of the manuscript. This work was supported by a grant from the European Space Agency, Prodex (BELSPO, Belgian Federal Government). BPD is a postdoctoral researcher of the Fonds de la Recherche Scientifique – FNRS (Belgium).

## References

Adams MJ, Johnson SA, Lefèvre P, Lévesque V, Hayward V, Adams MJ, Johnson SA, Lefe P, Thonnard JL, Hayward V, Andre T, André T, Thonnard JL, Levesque V. Finger pad friction and its role in grip and touch. J R Soc Interface 10: 20120467, 2013.

André T, Lefèvre P, Thonnard J. Fingertip moisture is optimally modulated during object manipulation. J Neurophysiol 103: 402–408, 2010.

André T, Lévesque V, Hayward V, Lefèvre P, Thonnard J. Effect of skin hydration on the dynamics of fingertip gripping contact. J R Soc Interface 8: 1574–1583, 2011.

Augurelle A-S, Smith AM, Lejeune T, Thonnard J-L. Importance of cutaneous feedback in maintaining a secure grip during manipulation of hand-held objects. J Neurophysiol 89: 665–671, 2003.

Barrea A, Cordova Bulens D, Lefevre P, Thonnard J-L. Simple and Reliable Method to Estimate the Fingertip Static Coefficient of Friction in Precision Grip. IEEE Trans Haptics 9: 492–498, 2016.

Barrea A, Delhaye BP, Lefèvre P, Thonnard J-L. Perception of partial slips under tangential loading of the fingertip. Sci Rep 8: 7032, 2018.

Bradski G. The OpenCV Library. Dr Dobb’s J Softw Tools, 2008.

Cadoret G, Smith AM. Friction, not texture, dictates grip forces used during object manipulation. [Online]. J Neurophysiol 75: 1963–9, 1996 http://www.ncbi.nlm.nih.gov/pubmed/8734595.

Delhaye B, Lefèvre P, Thonnard J-L. Dynamics of fingertip contact during the onset of tangential slip. J R Soc Interface 11: 20140698, 2014.

Delhaye BP, Barrea A, Edin BB, Lefèvre P, Thonnard J-L. Surface strain measurements of fingertip skin under shearing. J R Soc Interface 13: 20150874, 2016.

Delhaye BP, Jarocka E, Barrea A, Thonnard J-L, Edin B, Lefèvre P. High-resolution imaging of skin deformation shows that afferents from human fingertips signal slip onset. Elife 10, 2021.

Delhaye BP, Long KH, Bensmaia SJ. Neural Basis of Touch and Proprioception in Primate Cortex. In: Comprehensive Physiology. John Wiley & Sons, Inc., p. 1575–1602.

Flanagan JR, Wing AM. The role of internal models in motion planning and control: evidence from grip force adjustments during movements of hand-held loads. J Neurosci 17: 1519–28, 1997.

Goodwin AW, Wheat HE. Sensory signals in neural populations underlying tactile perception and manipulation. Annu Rev Neurosci 27: 53–77, 2004.

Gueorguiev D, Bochereau S, Mouraux A, Hayward V, Thonnard J-L. Touch uses frictional cues to discriminate flat materials. Sci Rep 6: 25553, 2016.

Jenmalm P, Birznieks I, Goodwin AW, Johansson RS. Influence of object shape on responses of human tactile afferents under conditions characteristic of manipulation. Eur J Neurosci 18: 164–176, 2003.

Johansson RS, Flanagan JR. Sensorimotor Control of Manipulation. In: Encyclopedia of Neuroscience. 2009a, p. 593–604.

Johansson RS, Flanagan JR. Coding and use of tactile signals from the fingertips in object manipulation tasks. Nat Rev Neurosci 10: 345–359, 2009b.

Johansson RS, Westling G. Roles of glabrous skin receptors and sensorimotor memory in automatic control of precision grip when lifting rougher or more slippery objects. Exp Brain Res 56: 550–564, 1984.

Lucas B, Kanade T. An iterative image registration technique with an application to stereo vision. [Online]. IJCAI 130: 121–130, 1981 http://flohauptic.googlecode.com/svn-history/r18/trunk/optic_flow/docs/articles/LK/Barker_unifying/lucas_bruce_d_1981_2.pdf [27 Nov. 2013].

Macefield VG, Johansson RS, Häger-Ross C, Johansson RS. Control of grip force during restraint of an object held between finger and thumb: Responses of cutaneous afferents from the digits. Exp brain Res 108: 155–171, 1996.

Nowak DA, Hermsdörfer J, Glasauer S, Philipp J, Meyer L, Mai N. The effects of digital anaesthesia on predictive grip force adjustments during vertical movements of a grasped object. Eur J Neurosci 14: 756–762, 2001.

Pasumarty SM, Johnson SA, Watson SA, Adams MJ. Friction of the human finger pad: Influence of moisture, occlusion and velocity. Tribol Lett 44: 117–137, 2011.

Peters RM, Hackeman E, Goldreich D. Diminutive digits discern delicate details: fingertip size and the sex difference in tactile spatial acuity. J Neurosci 29: 15756–61, 2009.

Sankaran A, Jain A, Vashisth T, Vatsa M, Singh R. Adaptive latent fingerprint segmentation using feature selection and random decision forest classification. Inf Fusion 34: 1–15, 2017.

Shi J, Tomasi C. Good features to track. In: Proceedings of IEEE Conference on Computer Vision and Pattern Recognition CVPR-94. IEEE Comput. Soc. Press, p. 593–600.

Tada M, Kanade T. An imaging system of incipient slip for modelling how human perceives slip of a fingertip. 26th Annu Int Conf IEEE Eng Med Biol Soc 3: 2045–2048, 2004.

Tada M, Mochimaru M, Kanade T. How does a fingertip slip? - Visualizing partial slippage for modeling of contact mechanics. In: Eurohaptics 2006. 2006, p. 2–7.

Tada M, Shibata T, Ogasawara T. Investigation of the touch processing model in human grasping based on the stick ratio within a fingertip contact interface. IEEE Int Conf Syst Man Cybern vol.5: 6, 2002.

Wang Q, Hayward V. In vivo biomechanics of the fingerpad skin under local tangential traction. J Biomech 40: 851–860, 2007.

Westling G, Johansson RS. Factors influencing the force control during precision grip. [Online]. Exp brain Res 53: 277–84, 1984 http://www.ncbi.nlm.nih.gov/pubmed/6705863.

Witney AG, Wing AM, Thonnard J-L, Smith AM. The cutaneous contribution to adaptive precision grip. Trends Neurosci 27: 637–643, 2004.

